# Melanoidins Suppress Cucumber and Tomato Diseases Incited by Pathogens that are Oomycetes, Bacteria and a Virus

**DOI:** 10.64898/2026.11.07.765025

**Authors:** Yigal Elad, Dalia Rav-David, Ludmila Tsechansky, Ellen R. Graber

## Abstract

We recently reported that solutions of the nitrogenous compounds melanoidins (M), that are formed during the final stages of the Maillard Reaction, are inducers of plant resistance (Elad et al. 2026). M promoted immunity and induced disease resistance through an effect on the transcriptome, primarily through the salicylic acid pathway. Here we tested the effect of M on plant diseases that are caused by Oomycetes, bacteria and a tobamovirus. Drench and spray of M suppressed *Pythium aphanidermatum* damping off in cucumber, cucumber downy mildew (*Pseudoperonospera cubensis*), tomato late blight (*Phytophthora infestans*), bacterial speck (*Pseudomonas syringae* pv*. tomato*), bacterial canker (*Clavibacter michiganensis* subsp. *michiganensis*) and the Tobamovirus Tomato brown rugose fruit virus (ToBRFV) in tomato. In various experiments control was achieved by concentrations of 0.03 and 0.1% and on older and younger leaves and even in long term experiments where the M was applied for ca 11 weeks. It can be concluded that the disease resistance inducer M can suppress soilborne and foliar diseases that are caused by Oomycetes, bacteria and a virus of the genus Tobamovirus, in addition to the fungal pathogen *Botrytis cinerea* reported earlier.

## Introduction

Melanoidins (M) are nitrogenous compounds that are formed during the final stages of the Maillard Reaction, originally described in early twentieth century (Maillard 1912). The chemical pathways involved in the Maillard Reaction were described by J.E. Hodge (1953). The variability in backbone composition of melanoidins and lack of a distinct molecular structure results in heterogeneity in the melanoidins that vary depending on the starting reactants and their ratios, ranging from amino acid-based to protein-based (melanoproteins) or polysaccharide-based compounds (Morales et al. 2012). Recently, we reported that M promoted cellular immunity and disease resistance against the fungal plant pathogen *Botrytis cinerea* (Elad et al. 2026). M was found to promote immunity and induce disease resistance through a swift and transient effect on the transcriptome, primarily through the salicylic acid (SA) pathway, as was demonstrated by transcriptomics and genetic defense mutants (Elad et al. 2026). In the present research we tested if M affects plant diseases that are caused by Oomycetes, bacteria and a virus.

The Oomycete *Pythium aphanidermatum* causes damping off in many plant species (Ben Yephet and Nelson 1999; Jaiswal et al. 2018) including cucumber (Al Sa’di et al. 2007; Elad and Chet 1987; Sivan et al. 1984). Induced resistance of *P. aphanidermatum* by *Pseudomonas chlororaphis* and *Bacillus subtilis* was reported in pepper (Nakkeeran et al. 2006). Benzothiadiazole (BTH) pretreatment of cucumber seeds resulted in reduced damping off in cucumber seedlings (Salari et al. 2025). Pretreatment of cucumber growth medium with biochar resulted in induced resistance to *P. aphanidermatum* in cucumber that was attributed both to chemical and microbiological changes in the medium (Jaiswal et al. 2018, 2019).

Cucurbits downy mildew is caused by the Oomycete *Pseudoperonospora cubensis* (Savory et al. 2011). It potentially causes severe epidemics in cucumber (Barnea et al. 2022; Shtienberg et al. 2010). Induced resistance in cucumber against *P. cubensis* was obtained with nutritional elements such as P, K, and Mg (Barnea et al. 2022) and silicon (Yu et al. 2010).

Late blight, caused by the Oomycete *Phytophthora solani*, is a major disease in tomato and potato (Frenkel et al. 2010). In tomato leaves it causes severe epidemics in greenhouse grown crops and in open fields (Palti and Rotem, 1973; Shtienberg et al. 2010). Control of tomato late blight by induced resistance was reported for an isolate of *Trichoderma harzianum* (Chowdappa et al. 2013), plant growth-promoting rhizobacteria (PGPR, Yan et al. 2002) and [beta]-Aminobutyric acid (BABA, Cohen et al. 1994).

Tomato bacterial speck, caused by the bacterial pathogen *Pseudomonas syringae* pv. *tomato*, is a gram-negative bacterium that epidemiologically is specific to this crop under field conditions (Falik et al. 1983; Preston, 2000). Reports on induced resistance against tomato bacterial speck are limited. Yoo et al. (2018) reported that an isolate of *Aspergillus terreus* that is a plant growth promoter which controls bacterial speck disease by inducing resistance in tomato plants. The endophytic bacteria *Bacillus pumilus* and *B. amyloliquefaciens* induce resistance against bacterial speck in tomato plants. Induced resistance in this case had a slight negative effect on plant growth parameters such as plant height and plant dry weight (Lanna-Filho et al. 2017).

*Clavibacter michiganensis* subsp. *michiganensis*, the causal agent of bacterial canker and wilt of tomato, is an important bacterial pathogen worldwide that also caused severe epidemics in Israel (Blank et al. 2016). It is a gram-positive bacterium that causes wilting, leaf necrosis, and characteristic small spots (Kleitman et al. 2008; Savidor et al. 2012). Among the inducers of resistance that were reported to reduce bacterial canker of tomato are acibenzolar-S-methyl (ASM) (Soylu et al. 2003), an isolate of a rhizobacterium *Pseudomonas sp.* (Takishita et al. 2018) and the plant growth promoting bacteria *Bacillus subtilis*, *Pseudomonas fluorescens* or *Azotobacter chroococcum* (Kolomiiets et al. 2019).

Tomato brown rugose fruit virus (ToBRFV) is a virus of the *Tobamovirus* genus that was first described in 2014 in Israel (Luria et al. 2017) and Jordan in 2015 (Salem et al. 2016). The main hosts of ToBRFV are tomato and peppers (Salem et al. 2023). The virus causes symptoms including mosaic and distortion of leaves and brown, wrinkly spots (rugose) on fruits and it spreads short distances by mechanical contact (Luria et al. 2017; Salem et al. 2016, 2023). *Pseudomonas fluorescens* and *Burkholderia gladioli* isolates sprayed on tomato plants enhanced defense enzyme activities, upregulated gene expression, and reduced symptom severity while their combination was superior to the treatments with the individual bacteria suspensions (Zohoursoleimani et al. 2025). The α-aminophosphonate ester Dufulin, an activator of the salicylic acid (SA) pathway, controlled ToBRFV in tomato plants (Wang et al. 2026).

In view of the ability of M to induce resistance against a fungal pathogen (Elad et al. 2026), we tested if the resistance inducer may also affect the above-mentioned pathogens on cucumber and tomato. The tests of M solutions were aimed at the suppression of the soilborne *P. aphanidermatum* and foliar *P. cubensis* Oomycetes pathogens of cucumber, and in tomato, the Oomycete *P. solani*, the bacterial pathogens *P. syringae* pv. *tomato* and *C. michiganensis* subsp. *michiganensis* and the *Tobamovirus* ToBRFV.

## Materials and methods

### Plants’ growth and maintenance

Plants were grown in either 1 L or 10 L pots containing a growth mixture of 7:3, v/v peat–tuff (sorted to ≤8 mm) referred-to below as ‘plants-growth medium’. Fertigation was applied 2 to 3 times daily via drip irrigation using 5:3:8 NPK fertilizer (120:30:150 mg L ¹ N:P:K; EC 2.2 dS m ¹), maintaining 25–50% drainage (Elad et al. 2010, 2012, 2026; Gupta et al. 2024; Mehari et al. 2015). The experiments with plants were held in a 20-mesh net house (April–October 2018–2024) or in a glasshouse with 25% shade netting and whitewash (Livnat, Tapazol, Israel) during the rest of the year. Day/night temperatures ranged from 18–33°C. All net house and glasshouse experiments were carried out at the Volcani Institute, Rishon LeZion, Israel (31°58′09″N, 34°48′02″E). The plants were maintained pest- and pathogen-free until treatments were applied (Elad et al. 2010, 2012, 2026; Gupta et al. 2024; Mehari et al. 2015).

#### Tomato

Tomato (*Solanum lycopersicum*) cv. Ikram (Syngenta, Israel) seedlings were obtained from Hishtil Nurseries (Israel) at age of 4 to 5 weeks post sowing. The plants were transplanted to pots and grown for 4–6 additional weeks before being exposed to pathogens. The tomato plants that were used in short term experiments were grown in 1 L pots and the plants used in commercial-like long-term experiments were grown in 10 L containers (Elad et al. 2026; Gupta et al. 2022a,b, 2024). In both cases there was one tomato plant in each pot, and it was held vertically by clipping it with plastic rings to bamboo sticks.

#### Cucumber

Cucumber (*Cucumis sativus*) cv. Manny (Genesis Seeds, Israel) was grown in 1 L pots. Cucumber plants were used as seedlings for *Pythium aphanidermatum* assays. The cucumber seeds were seeded in pots containing pathogen free growth medium (5 seeds per pot), and one day after emergence the plants were transplanted into infested medium as described below (Jaiswal et al. 2018; Philosoph et al. 2018).

### Plant diseases inoculation and evaluation

Cucumber damping-off was incited by using millet seeds inoculum of *Pythium aphanidermatum*. For this purpose, *P. aphanidermatum* was cultured on PDA and grown at 25°C. Mycelium discs were transferred to autoclaved millet in 250 ml Erlenmeyer flasks, and the mixture was incubated for 5 days. Once the mycelium occupied the seeds it was homogenized and mixed into the plants-growth medium at a concentration of 0.25% (w/w). Seedlings were transplanted into the infested plants-growth medium, and mortality was recorded every 2 days. Typical death of seedlings was observed and documented as damping off mortality (Jaiswal et al. 2018; Philosoph et al. 2018).

*P. apanidermatum* linear growth was recorded in PDA Petri plates. Mycelium discs from 3 days old cultures was inoculated on PDA plates amended with M in concentrations of 0.01, 0.03, and 0.1%. The plates were incubated at 25°C and the diameter of colonies was recorded one and two days later. The area of the colony of the oomycete was calculated.

Cucurbits downy mildew (*Pseudoperonospora cubensis*) developed naturally on 1 to two months old cucumber plants during fall, winter and spring (November to April) under net-house conditions and was evaluated for severity as leaf coverage (0–100%) with typical downy mildew symptoms (Barnea et al. 2022).

Tomato late blight (*Phytophthora solani*) developed naturally during fall, winter and spring (November to April) under net-house and greenhouse conditions and was evaluated for severity as coverage (0–100%) with typical late blight symptoms (Frenkel et al. 2010; Shtienberg et al. 2010).

Tomato bacterial speck (*Pseudomonas syringae* pv. *tomato*) developed naturally during fall, winter and spring (November to April) under net-house and greenhouse conditions and was evaluated for severity as coverage (0–100%) with typical bacterial speck symptoms (Bashan et al. 2002; Falik et al. 1983).

Bacterial canker (*Clavibacter michiganensis* subsp. *michiganensis*) was cultured on LB agar at 28°C, suspended in water (5 × 10 CFU/mL), and sprayed onto tomato plants (Savidor et al. 2012; Verma et al. 2026). Plants were maintained at 25 ± 2°C, and disease severity was rated on a 0– 100% scale.

Tomato brown rugose fruit virus (ToBRFV) was inoculated artificially. Israeli isolate of ToBRFV, ToBRFV-IL (KX619418.1) was used for ToBRFV infection. Tomato plants were sprayed with carborundum powder suspension prior to inoculation, the plants were let to passively dry and then the leaves were rubbed with a phosphate buffer solution (0.01 M, pH 7.0) supplemented with crushed leaves from the ToBRFV-infected tomato plants. The severity of ToBRFV symptoms were evaluated on a scale of 0-100% (Alon et al. 2021; Klap et al. 2020). RT-PCR analysis was carried out according to Alon et al. (2021).

### M synthesis and application

The melanoidins were synthesized by means of Maillard reaction in a dry synthesis method at 150°C using L-glutamic acid and D-glucose. Powders of the ingredients were mixed with each other at an equimolar ratio and ground together as described recently (Elad et al. 2026). The M powder was ground, placed in hermetically sealed vials, and kept in a desiccator with silica gel at room temperature until used (Elad et al. 2026). Plants were sprayed or drenched with M water solutions at concentrations of 0.03 and 0.1%. Spray applications were performed using a 1 L hand sprayer calibrated to emission of fine droplets until full canopy coverage was achieved. The solution was applied at volume of 8–15 mL per each mature plant depending on plant size. Drench applications consisted of 10–30 mL per pot, covering the volume of root zone with ∼5% drainage (Elad et al. 2026).

The control of *P. aphanidermatum* damping off by M was compared with a common fungicide. The medium was drenched once with a fungicide, 0.25% Dynon in water drench, containing 722 g/L Propamocarb HCl, Bayer, Germany. The inducer of resistance monopotassium phosphate (MKP; KH PO , ICL Haifa, Israel) was used in one experimental set (Reuveni et al. 1993; Elad et al. 2021).

### Calculations and statistics

Percentage data were arcsine-transformed prior to analysis. Area under the disease progress curve (AUDPC) or area under mortality progress curve (AUMPC) were calculated where applicable. Standard errors (SE) are presented as bars in the figures. Replication ranged from 5 to 6 per treatment. Data (disease severity, AUDPC, and AUMPC) were analyzed by ANOVA followed by Tukey–Kramer HSD test, using JMP software (SAS Institute, Cary, NC, USA; α = 0.05). Different letters indicate significant differences among treatments.

Disease reduction (DR, %) was calculated as:

DR = 100 – 100 × (Tn / Tcon), where Tn is the disease level in treated plants, and Tcon is the untreated control. DR was calculated for each replicate and averaged for each treatment.

## Results

### Effect of M on a soilborne Oomycete – *Pythium aphanidermatum* damping off in cucumber seedlings

*P. aphanidermatum* damping off in the untreated control reached 41.25% incidence (Fig. S1). Spray of M at concentrations of 0.03 and 0.1% significantly reduced *P. aphanidermatum* damping off during days 10 and 17; at two of the dates the 0.1% M was significantly better than 0.03% M in reducing damping off (Fig. 1a). The results in progress curves presented in Fig. S1 were calculated in order to analyze the area under mortality progress curve (AUMPC). The AUMPC was significantly reduced by 52.2 and 62.7% by 0.03 and 0.1%, respectively (Fig. 1A). *P. aphanidermatum* damping off reached 45% in the untreated control 18 days after the initiation of the M treatments that are summarized ni Fig. 1b. M drench and spray reduced *P. aphanidermatum* damping off in young cucumber plants by 67.7 and 76.0%, respectively, and dynone (propamocarb) reduced it by 97.0% (Fig. 1b).

**Fig. 1.**
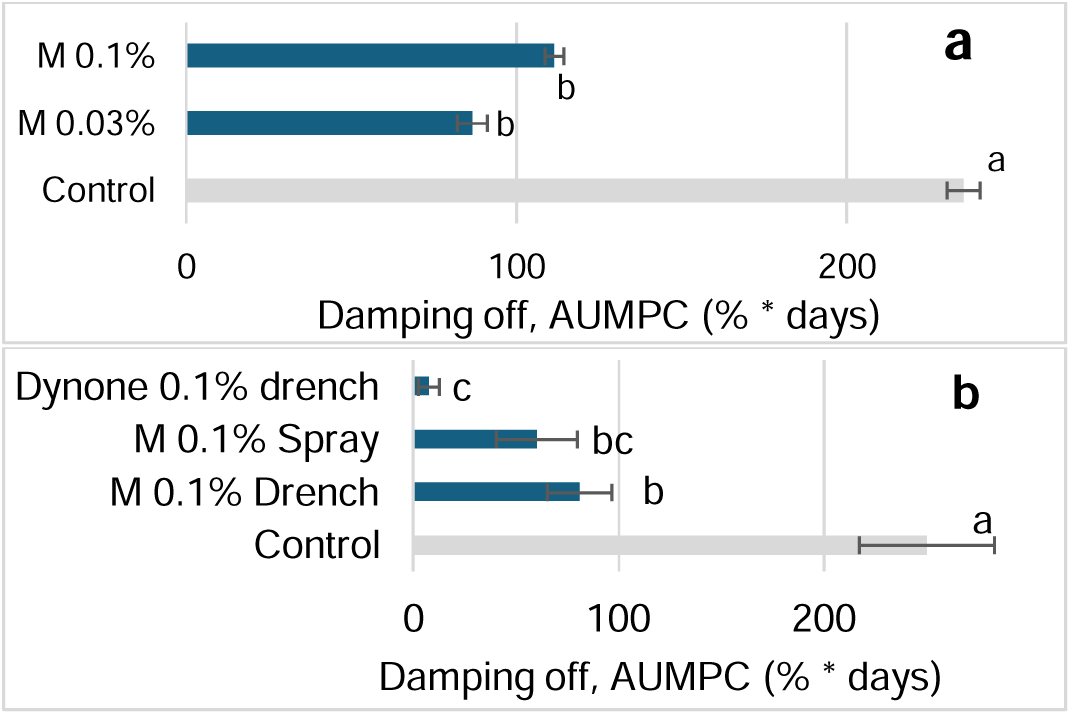
Effect of M on Oomycetes incited soilborne disease in cucumber – Pythium damping off. **a**. M spray was applied at concentrations of 0.03 and 0.1% on the day of seedlings transplanting in the *Pythium aphanidermatum* amended growth mixture and 7 days afterwards as described in Fig. S1 and the area under the mortality progress curve (AUMPC) of data presented in Fig. S1 was calculated through 17 days after transplanting. **b**. M was applied by spraying and by drenching and the chemical pesticide, dynone (Propamocarb HCl, Bayer) was drenched in cucumber seedlings to control the mortality of cucumber seedlings due to *P. aphanidermatum* in a contaminated growing medium. M was applied at 0, 7, and 14 days after seeding and dynone was applied at 0 and 7 days. (AUMPC) through 18 days after seeding. Damping off was evaluated every 2 to 3 days and presented as percent of all plants in each pot. Percent data at each evaluation date (Fig. S1) and AUMPC values in each graph followed by a different letter are significantly different according to one-way ANOVA with Tukey’s HSD test. Default significance levels were set at α = 0.05. Bars = SEs.

### Effect of M on the growth of *Pythium aphanidermatum* in culture

The M was mixed in PDA growth medium in concentrations of 0.01, 0.03, and 0.1%, and the pathogen’s mycelium growth on it was followed for 2 to 3 days. After two days *P. aphanidermatum* reached 41.8 cm^2^ in the non-amended medium and 37.4 to 44.2 cm^2^ in the amended media with no significant difference between treatments.

### Effect of M on a foliar Oomycetes pathogen – *Pseudoperonospera cubensis* downy mildew (PcCDM) in cucumber

In a short-term experiment, we tested if 0.1% M applied by spray and drench affects *P. cubensis* downy mildew (PcCDM) on cucumber leaves (Fig. S2). M treatments significantly reduced the severity of PcCDM by 70.8 to 73.8%, respectively (Fig. S2). M spray and drench treatments of 0.03 and 0.1% were applied under commercial-like conditions on a weekly basis. PcCDM was significantly reduced by 41.1-61.3, 49.1-61.4 by the four treatments at 41, 58, and 71 days after treatments’ initiation, respectively, while at 71 days only the 0.1% M drench and spray treatments significantly reduced the disease severity, achieving 51.8-53.1% reduction (Fig. 2a). In a second commercial-like setup, M 0.1% spray and drench treatments were compared with the Mono potassium phosphate (MKP) disease resistance inducer. PcCDM evaluated at 61 days after treatments initiation was 41.2 and 16.8% on leaves 8 and 13, respectively (Fig. 2B). The disease severity was reduced by 60.6 and 76.1, 93.9, and 100 and 91.5 and 100% the MKP, and M spray and drench treatments on the respective leaves, respectively (Fig. 2b). The extent of efficacy on the two leaves was similar.

**Fig. 2.**
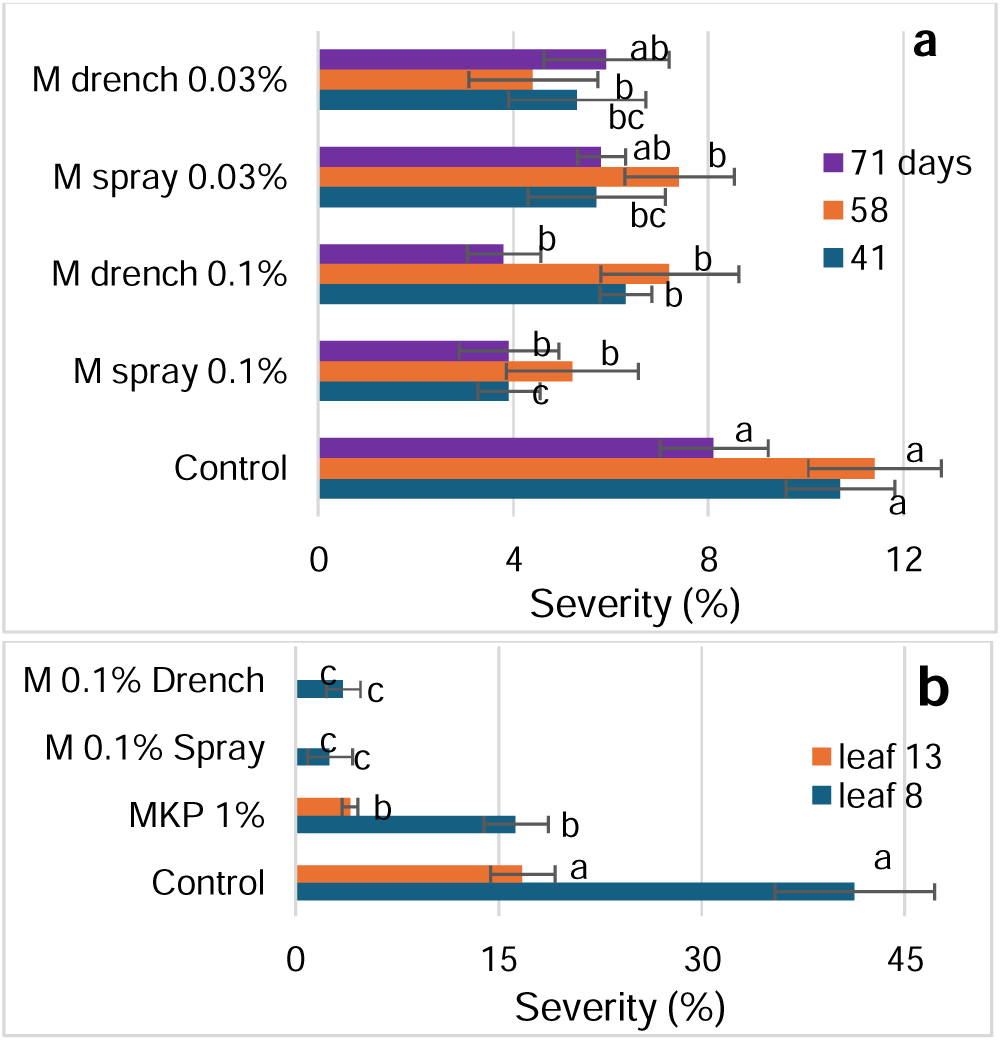
Effect of M on an Oomycetes incited foliar disease in cucumber – Control of *Pseudoperonospora cubensis* cucumber downy mildew (PcCDM). **a**. Control of naturally developed PcCDM by 0.03 and 0.1% M spray and drench in a commercial-like set up where applications were carried out on a weekly basis and disease severity was evaluated at 41, 58, and 71 days after treatments initiation. **b**. Control of naturally developed PcCDM by 0.1% M drench and spray and by MKP (KH_2_PO_4_, ICL Haifa, Israel) spray, applied on a weekly basis. PcCDM was evaluated on two leaves at 61 days after sprays initiation. Severity of PcCDM was evaluated on a 0 to 100% scale, in which 0 = healthy leaves and 100% = leaves completely covered by the damage symptoms. Values in each date (A) and each leaf (B) followed by a different letter are significantly different according to one-way ANOVA with Tukey’s HSD test. Default significance levels were set at α = 0.05. Bars = SEs.

### Tomato late blight (*Phytophthora infestans*) affected by M

Late blight (*P. infestans*) occurred naturally on tomato plants and reached in the untreated control, 19.4% severity 13 days after the initial M treatment. The M 0.1% spray that was applied twice (at days 0 and 5) significantly reduced disease severity on stem and leaves at all evaluation dates (Fig. S3); It significantly reduced the severity as calculated for AUDPC through days 1 to 13 by 51.4% (Fig. 3a). One application of 0.03% M spray reduced the tomato late blight by 57.3% (Fig. 1b). Late blight on tomato plants in an experiment that was carried out under commercial-like conditions was treated weekly with 0.03 and 0.1% M drenches and sprays. The disease was significantly reduced through 76 days after initiation at the 0.1% spray treatment by 46.1 to 76.0%; at the 0.1% drench by 63.6 to 100%; at the 0.03% spray by 75.8 to 94.7%. The 0.03% drench significantly reduced the disease by 57.4 to 86.7% during days 33 to 62 after treatments initiation (Fig. 3c). The calculated AUDPC was significantly reduced by 51.9, 74.3, 85.5, and 58.3% by M 0.1% spray and drench and by 0.03% M spray and drench, respectively (Fig. 3d).

**Fig. 3.**
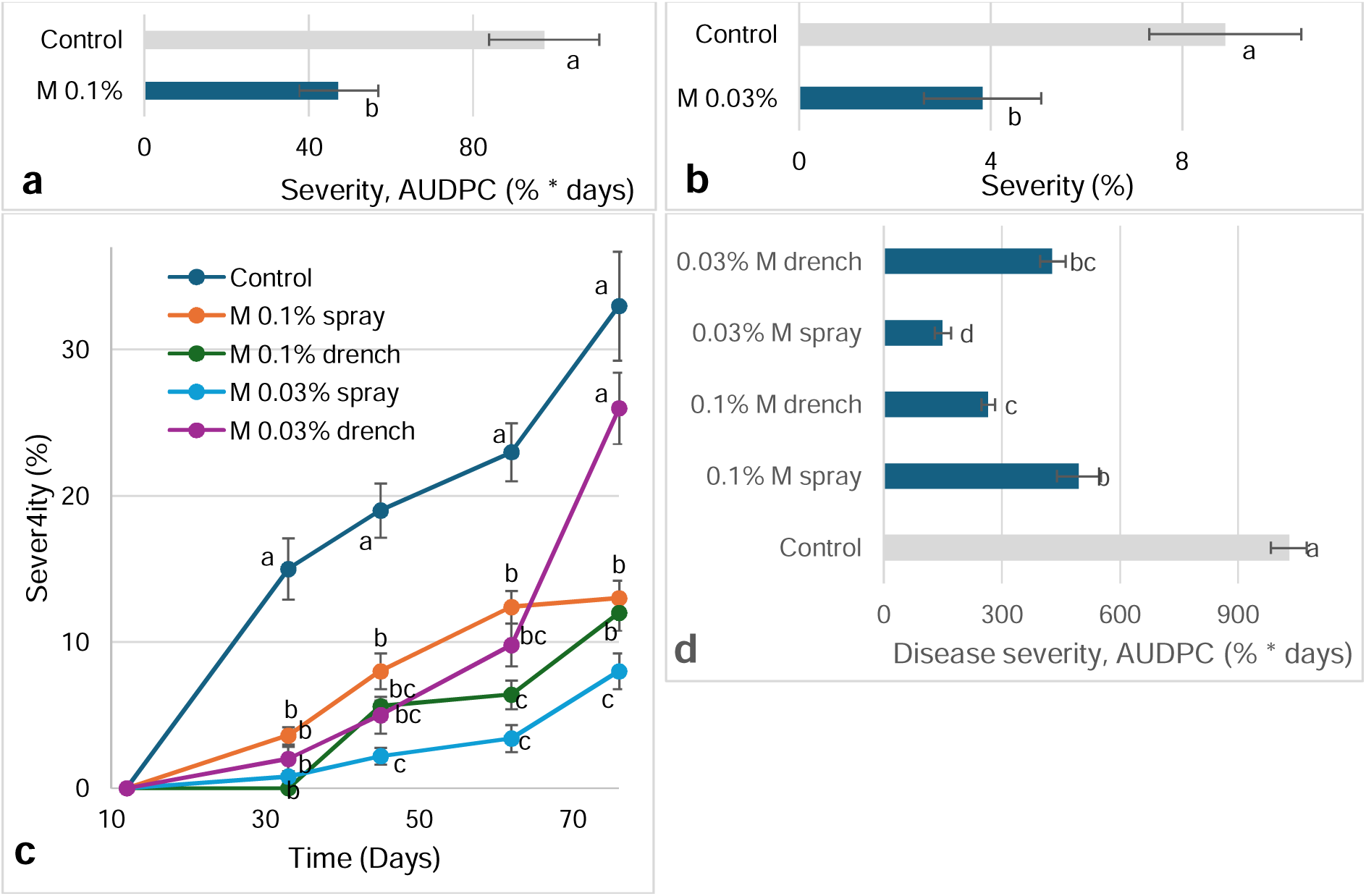
Effect of M on late blight caused by *Phytophthora infestans* in tomato plants. **a**. Plants were treated by 0.1% M spray at days 0 and 5 and disease severity was evaluated until day 13 after the initiation of M applications as described in Fig. S3 and area under disease progress curve (AUDPC) was calculated. **b**. Plants were treated by 0.03% M spray once and disease severity was evaluated at day 5 after the initiation of M applications. **c**. Tomato plants were treated 11 times on a weekly basis by M spray and drench at concentrations of 0.03 and 0.1% and disease severity was evaluated 4 times until day 76 after treatments initiated. **d**. AUDPC was calculated for the data on treatments described in **c**. Severity of late blight was evaluated on a 0 to 100% scale, in which 0 = healthy leaves and 100% = leaves and stems completely covered by the damage symptoms and AUDPC was calculated as mentioned above. Values in each date and each graph followed by a different letter are significantly different according to one-way ANOVA with Tukey’s HSD test. Default significance levels were set at α = 0.05. Bars = SEs.

### M controls bacterial speck (*Pseudomonas syringae* pv*. tomato*) on tomato canopy

Bacterial speck (*P. syringae* pv*. tomato*, PstBS) occurred naturally on the tomato plants. The disease was suppressed with 0.1% M drench and spray applications at 11 and 13 days in a similar manner on lower and upper leaves (Fig. 4a). Disease severity in the untreated control reached 52.5% in an experimental set in which the M was treated with 0.03% concentration (Fig. S4). In this experiment the severity of disease was significantly reduced at dates 6 through 13 by the 0.03% M drench and spray treatments (Fig. S4). Disease severity, as calculated for AUDPC, was reduced by 77.5-80.0% and 51.1-61.9 by the M concentrations 0.1% (Fig. 2b) and 0.03% (Fig. 2c), respectively.

In another experimental setup PstBS severity was first evaluated 10 days after the first application of M (spraying of 0.1% M) and it was evaluated until 11 days later, scoring 37.3% severity on the whole plants in the untreated control. The M spray treatment reduced the disease by 49.4, 40.5, 72.0, and 51.1% on the lower, middle, and upper leaves and the whole tomato plants, respectively (Fig. 2D), revealing a better disease reduction on the upper leaves where disease in the untreated control was lower than on the middle and upper leaves (21.4, 32.6, and 57.9%, respectively in upper, middle and lower leaves).

**Fig. 4.**
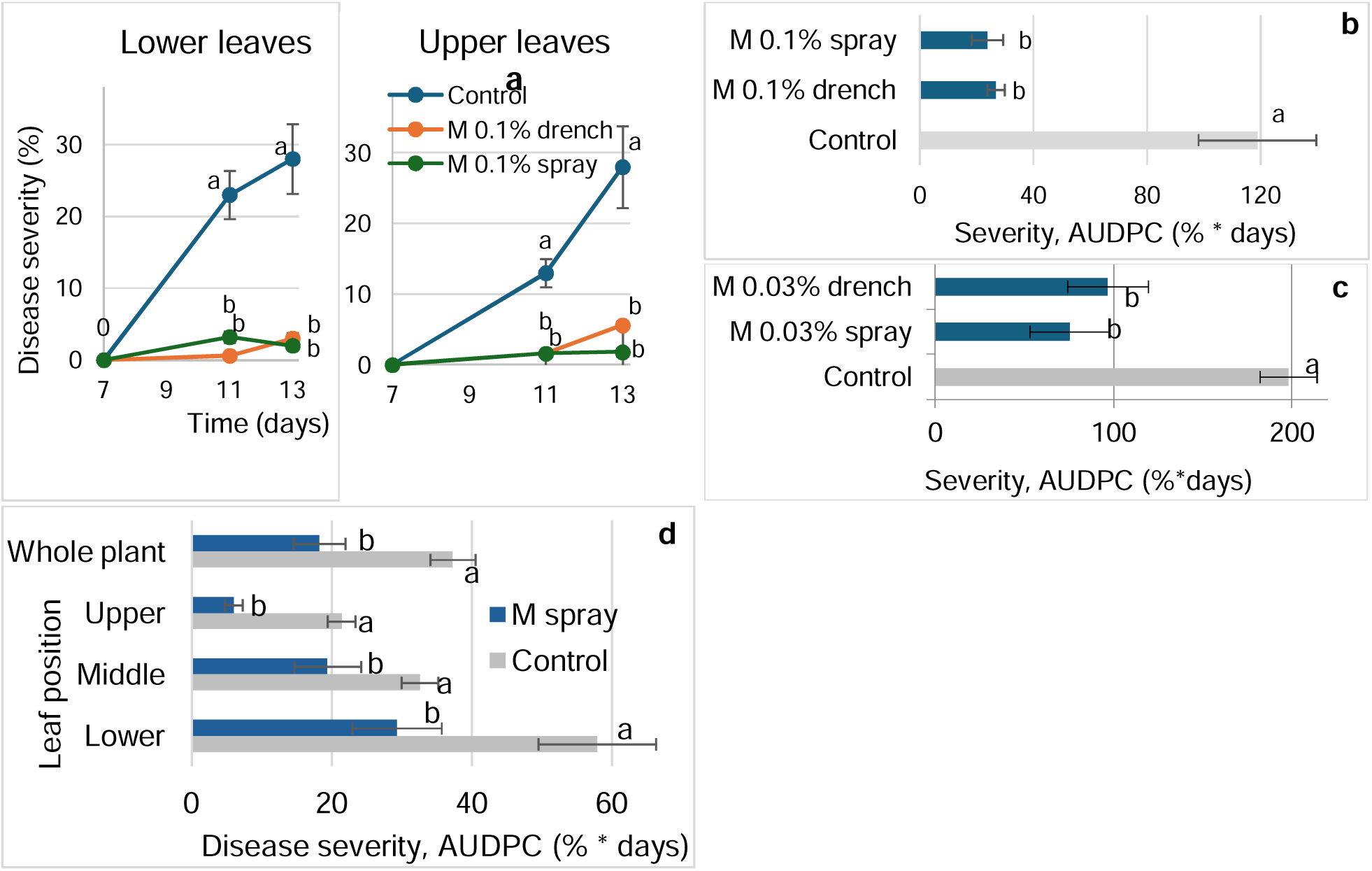
Effect of M on the severity of *Pseudomonas syringae* pv*. tomato* bacterial speck (PstBS) naturally developed on leaves of tomato plants. **a**-**c**. Effect of M spray and drench applied at 0 and 4 days after experiment initiation at concentrations of 0.1% (**a,b**) and 0.03% (**c**) and evaluated through 13 days. Disease was evaluated on lower and upper leaves at two dates (**a**) and are also presented as area under disease progress curve (AUDPC) for whole plants treated by 0.1% M and evaluated for PstBS through 13 d (**b**, details described in Fig. 4a), and AUDPC of disease severity through 13 days in untreated plants and plants treated by M 0.03% drench and spray (**c**, detailed results are in Fig. S4). **d**. Effect of 0.1% M applied by spraying at 0, 7 and 14 days on the severity of PstBS on lower, middle, and upper leaves and on whole plants (averaged data of the 3 leaf levels=whole plant) as evaluated through 21 days and calculated for AUDPC. Severity of PstBS was evaluated on a 0 to 100% scale, in which 0 = healthy leaves and 100% = leaves completely covered by the damage symptoms. AUDPC was calculated through 13 days (**b** and **c**) and through 21 days (**d**). Values in each date and leaf height (**a**), graph (**b**,**c**) and in each leaf level (**d**) followed by a different letter are significantly different according to one-way ANOVA with Tukey’s HSD test. Default significance levels were set at α = 0.05. Bars = SEs.

### Effect of M on *Clavibacter michiganensis* subsp. *michiganensis* bacterial canker (CmmBC) in tomato

The bacterial canker disease developed after artificial inoculation with suspension of *C. michiganensis* subsp. *michiganensis*. Typical symptoms that developed on the leaves consisted of small necrotic dots that spread over the leaflets area (Fig. 5a) and dry rot that spread on the leaflets’ edges (Fig. 5b). The small dots’ symptoms were not affected by the M treatments at the early evaluation of 8 days, and it was significantly reduced on day 11 by 48.9 and 38.5% by the 0.1% M drench and spray, respectfully (Fig. 5a). The edges dry rot evaluated on day 8 was only 37.5% reduced by the M spray while on day 11 both treatments significantly reduced their severity by 63.0 and 60.1%, respectfully (Fig. 5b).

**Fig. 5.**
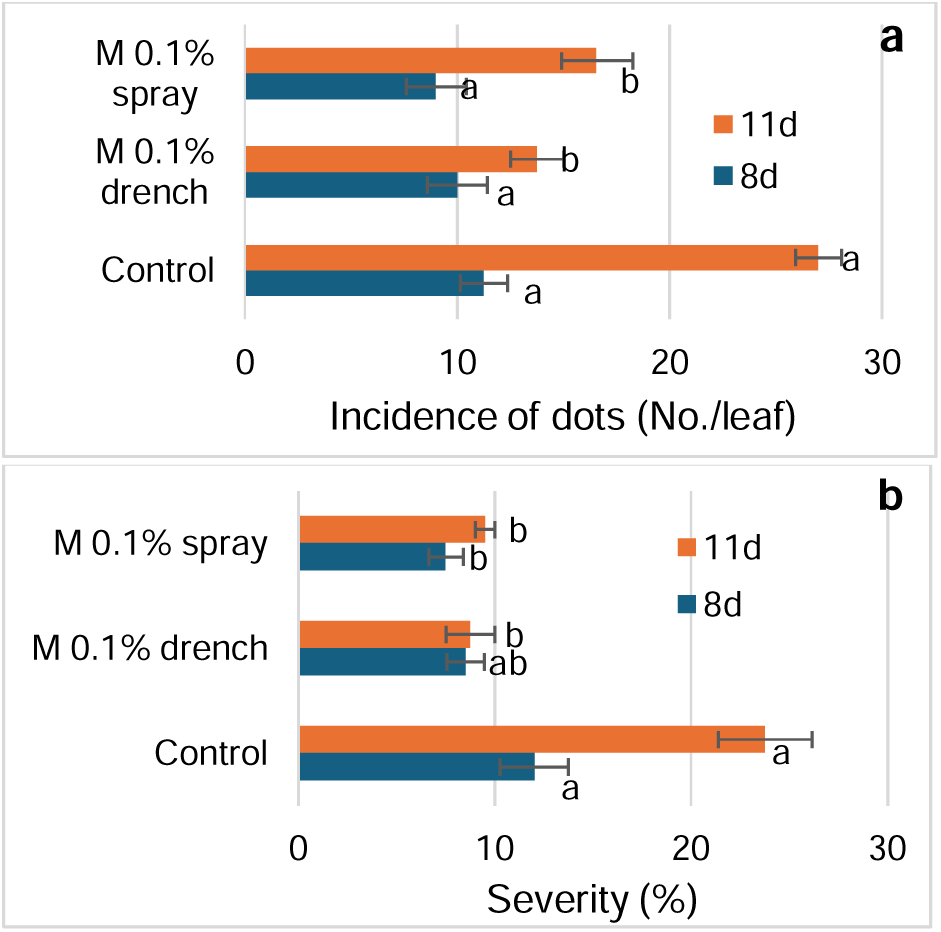
Effect of M 0.1% drench and spray applied at days 0 and 3 on the severity of *Clavibacter michiganensis* subsp. *michiganensis* bacterial canker (CmmBC) on tomato leaves. Bacterial suspension was used at day 0 to artificially infect the plants. Symptoms consisted of dots that were spread on the leaf blade (**a**) and spreading rot that developed on leaflet edges (**b**). Severity of each of the typical symptoms was evaluated at two dates (days 8 and 11) on a 0 to 100% scale, in which 0 = healthy leaves and 100% = leaflets completely covered by the damage symptoms. Values in each date and symptom type followed by a different letter are significantly different according to one-way ANOVA with Tukey’s HSD test. Default significance levels were set at α = 0.05. Bars = SEs.

### M affects Tomato brown rugose fruit virus (ToBRFV)

Tomato brown rugose fruit (ToBRFV), Tobamovirus, is transmitted through mechanical contact and that was the way of its introduction to the tomato plants in our experiments. Treatments of 0.1% spray and drench were carried out 3 days before infection, on the infection day, and 8 days later. The severity of the ToBRFV symptoms in tomato plants of the untreated control reached 36.0%, 25 days after infection (Fig. 6**a**). Disease symptoms were reduced significantly and similarly until day 21 after treatments initiation while at day 25 the severity reduction was 63.9 and 33.3% in the drench and spray treatments, respectively, and the drench treatment was significantly superior to the spray treatment (Fig. 6**a**). The AUDPC calculation revealed 37.2 and 62.5% reduction, respectively, in the spray and drench treatments (Fig. 3**b**).

The level of ToBRFV presence in samples of leaves taken in the experiment treatments was 60, 40, and 20% in the untreated control, M spray and M drench, respectively, revealing an effect of inhibition of virus presence and multiplication in the leaves by M treatment.

**Fig. 6.**
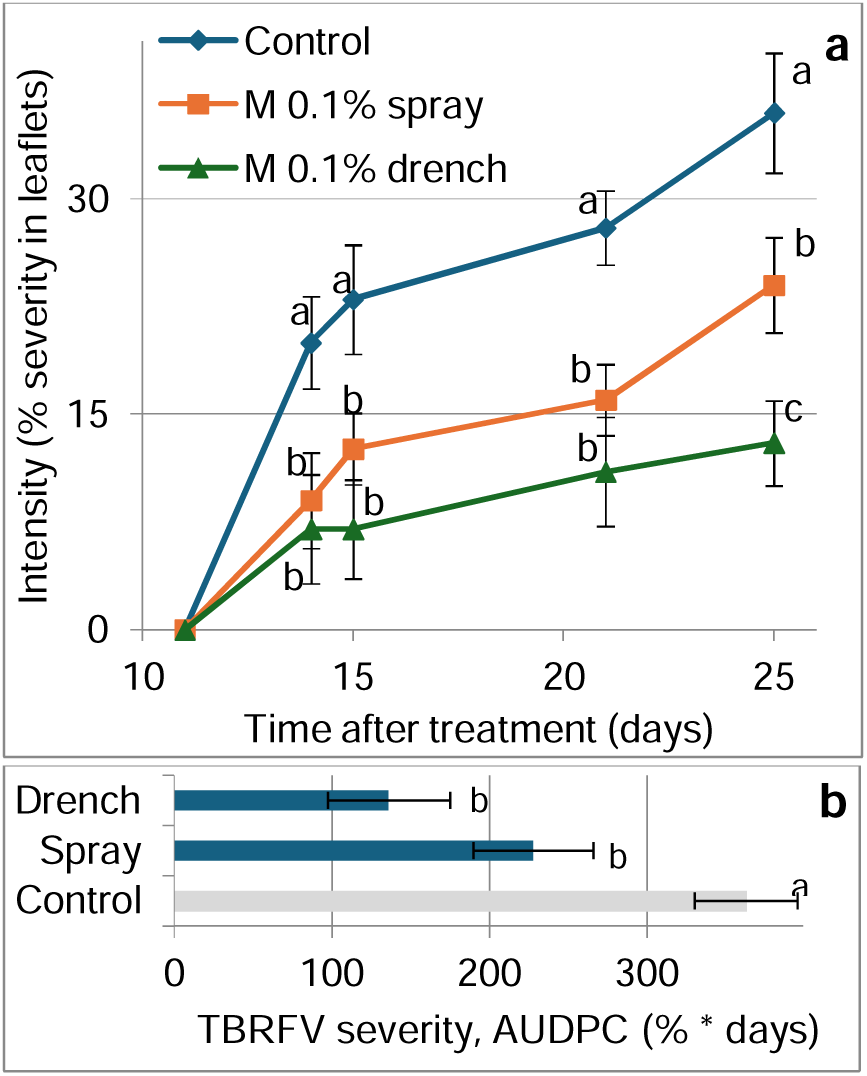
Effect of M on the tomato viral disease Tomato brown rugose fruit Tobamovirus (ToBRFV). M (0.1%) spray and drench treatments were carried out 3 days before infection, on the infection day, and 8 days later. Infection was carried artificially and the ToBRFV symptoms on the leaves were evaluated through 25 days after infection. The severity of symptoms on a 0 to 100% scale, in which 0 = healthy leaves and 100% = leaves completely covered by the viral symptoms. AUDPC was calculated through days 10 to 25. Values in each date (**a**) and graph (**b**) followed by a different letter are significantly different according to one-way ANOVA with Tukey’s HSD test. Default significance levels were set at α = 0.05. Bars = SEs.

## Discussion

We recently reported that M solutions affected the severity of the necrotrophic fungal plant pathogen *Botrytis cinerea*, the causal of gray mold in tomato (Elad et al. 2026). Here we demonstrate that M also suppresses diseases that are caused by pathogens that do not belong to the fungal kingdom. The pathogens that were controlled are the Oomycetes *Py. aphanidermatum* that causes damping off in cucumber (Figs 1 and S1), foliar diseases caused by Oomycetes downy mildew in cucumber (*Ps. Cubensis*, Figs 2 and S2) and late blight in tomato (*Ph. infestans*, Figs 3 and S3). In tomato M also controlled the bacterial diseases bacterial speck caused by *Ps. syringae* pv*. tomato* (Figs 4 and S4) and bacterial canker caused by *C. michiganensis* subsp. *michiganensis* (Fig. 5). Interestingly, the viral disease ToBRFV was also affected by M in tomato plants (Fig. 6).

M is not toxic to the pathogens as was shown here for *Py. aphanidermatum* in a culture medium and shown earlier for *B. cinerea* (Elad et al., 2026). We reported that M activity is due to promotion of cellular immunity and by induction of disease resistance against *B. cinerea* (Elad et al. 2026). M promotes immunity and induces disease resistance through a swift and transient effect on the transcriptome, primarily through the salicylic acid (SA) pathway, as was demonstrated by transcriptomics and genetic defense mutants (Elad et al. 2026).

Induced resistance is commonly divided into induced systemic resistance (ISR) and systemic acquired resistance (SAR) (Walters and Heil 2007). SAR biosynthesis pathway generally relies on SA, which has been shown to increase in the plant tissues, during pathogenesis and involves the activation of pathogenesis-related (PR) genes (Yalpani et al. 1991). SA (as well as jasmonic acid that is related to ISR) (Gupta et al. 2023; Pieterse et al. 1998; Thulke and Conrath, 1998), SA analogs such as benzothiadiazole (BTH) (Leibman-Markus et al. 2023; Meller Harel et al. 2014; Perazzolli et al. 2008), and several amino acids including phenylalanine, glutamine (Gupta et al. 2020; Liao et al. 2024; Oliva et al. 2020), are among the reported inducers of plant immunity. Systemic activity of M was demonstrated here in suppression cases of *Py. aphanidermatium* damping off (Fig. 1B), *Ps. cubensis* cucurbits downy mildew (Figs 2 and S2), *Ph. infestans* late blight (Fig. 3 c,d), *Ps. syringae* pv*. tomato* bacterial speck (Figs 4a-c and S4), *C. michiganensis* subsp. *michiganensis* bacterial canker (Fig. 5) and ToBRFV (Fig. 6).

M induced resistance not only against foliar diseases but also against a soilborne disease that causes damping off in cucumber (Figs 1 and S1) as was also shown in pepper seedlings with two biocontrol bacteria that induce resistance (Nakkeeran et al. 2006), BTH pretreatment of cucumber seeds (Salari et al. 2025) and pretreatment with biochar (Jaiswal et al. 2018, 2019). BTH is a SAR inducer that activates the SAR signal transduction pathway at the site of or downstream of SA accumulation (Friedrich et al. 1996) while also M induces resistance through the SA pathway (Elad et al. 2026).

In the case of *Py. aphanidermatum* damping off, we compared M with a common chemical, dynone (propamocarb) that statistically was like the spray treatment, but was significantly better than the drench treatment (Fig. 1). MKP is an inducer of resistance that was tested in cucumber crops (Reuveni et al. 1993; Elad et al. 2021). In the present study, cucumber downy mildew was suppressed better by M than by MKP on leaves of two ages at 61 days after initiation of the applications (Fig. 2b).

Comparison of drench versus spray application methods and two concentrations, 0.03 vs. 0.1%, revealed that both application methds and concentrations were similarly effective in most cases. The exceptions were late blight, where the results were inconsistent (Fig. 3d), and the last evaluation date (day 25) of ToBRFV, where the spray application was significantly less effective than the drench (Fig. 6a). Yet, overall, there was no significant difference in the calculated AUDPC between drench and spray effect for ToBRFV (Fig. 6b).

ToBRFV is a Tobamovirus that is transmitted by mechanical contact (Luria et al. 2017; Salem et al. 2016, 2023). There are no reported control agents for it, and few effective resistance inducers. Resistance inducers for ToBRFV include sprayed isolates of *Ps. fluorescens* - and *Burkholderia gladioli* on tomato plants (Zohoursoleimani et al. 2025) and the activator of the SA pathway, α-aminophosphonate ester Dufulin (Wang et al. 2026). Here we showed that M also has this activity (Fig. 6).

M significantly reduced bacterial speck better on younger leaves than on medium age and older leaves (Fig. 4d). Better activity in the younger leaves may be because of longer exposure of their tissue to the signals of induced resistance than the exposure of the leaf tissue to the pathogen infection and disease development while in older leaves the exposure to induced resistance signals is similar to the duration of exposure to the pathogen. Interestingly, the induction of resistance by M occurred also in experiments that were carried out under commercial-like conditions and lasted even for 9 to 11 weeks of repeated applications as was demonstrated for downy mildew in cucumber (Fig. 2 A) and late blight in tomato (Fig. 3c,d).

Inducers of resistance in various plant species – pathogen types are chemicals such as nutritional elements (Gupta et al. 2022b), BABA (Cohen et al. 1994), Dufulin (Wang et al. 2026) and ASM (Meller Harel, 2014; Perazzoli et al. 2008), fungi and bacteria such as biocontrol control agents (Gupta et al. 2022b, 2024; Meller Harel, 2014; Perazzoli et al. 2008) and PGPRs (Yan et al. 2002), and physical treatments like root zone warming (Gupta et al. 2021). We report here the induction of resistance by a chemical inducer, M, against the various Oomycetes, bacterial and viral pathogens in cucumber and tomato plants. Reports in the literature regarding the type of inducers of the diseases that are reported here as targets of M vary with the specific pathogen, as follows. Similarly to M as chemical inducer, induction of resistance towards *Py. aphanidermatum* damping off was achieved with inducers of chemical nature such as biochar (Jaiswal et al. 2018, 2019) and BTH (Salari et al. 2025); *Ps. cubensis* downy mildew was suppressed with chemical inducer such as nutritional elements (Barnea et al. 2022) and silicon (Yu et al. 2010); Induction of resistance against *Py. solani* late blight was achieved with the chemical BABA (Cohen et al. 1994); We did not identify chemical inducing resistance for Ps. *syringae* pv. *tomato* bacterial speck but the chemical inducer ASM was reported for *C. michiganensis* subsp. *michiganensis* bacterial canker; and the chemical Dufulin induced resistance against ToBRFV in tomato plants (Wang et al. 2026). All these cases of chemicals, including M, involve induced resistance that primarily is active through the SA pathway (Cohen et al. 1994; Elad et al. 2026; Gupta et al. 2022b; Meller Harel, 2014; Wang et al. 2026).

In conclusion, M treatments to tomato and cucumber plants resulted in suppression of diseases caused by Oomycetes, bacteria and a virus. M was found to act as a robust systemic inducer of SAR that leverages the host’s innate priming mechanisms to provide a broad-spectrum resistance. M is easily produced in a simple synthesis method, and it has high water solubility, so solutions of this resistance inducer are ideal for environmentally friendly disease management. M application by spray to the plant’s canopy and by drenching the roots were both effective in controlling foliar and soil-borne pathogens. M operates not through direct antimicrobial toxicity through the systemic activation of the host’s internal defense machinery (Elad et al. 2026).

## Authors Contributions

**Yigal Elad**: Writing – review & editing, Writing – original draft, Supervision, Methodology, Investigation, Formal analysis, Conceptualization. **Ellen R. Graber**: Writing – review & editing, Supervision, Methodology, Investigation, Funding acquisition, Formal analysis, Conceptualization. **Dalia Rav-David**: Methodology, Investigation, Formal analysis. **Ludmila Tsechansky**: Methodology, Investigation.

## Acknowledgments

We thank the Israel Innovation Authority and Adama Agricultural Solutions Ltd. Israel, for partial support of this work, and members of the Elad and Graber research groups for continuous support. We thank Dr. Omer Frenkel, Dept of Plant Pathology – Volcani Institute for support, advice and supply of *Py. aphanidermatum* inoculum, Dr. Daron Teper Dept of Plant Pathology – Volcani Institute for supplying *C. michiganensis* subsp. *michiganensis* suspension, Prof. Maya Bar Department of Life Sciences, Ben-Gurion University of the Negev, Beer-Sheva for long term assistance and advice, and Mr. Menachem Borenshtein, and Mr. Eyal Glantz, Dept of Plant Pathology for the technical support in the greenhouses.

## Data Availability Statement

The authors declare that the data supporting the findings of this study are available within the paper and its Supporting Information files. Raw data is available from the corresponding author upon reasonable request.

Research involving human participants and/or animals This article does not contain any studies with human participants or animals performed by any of the authors.

