## Supplemental Figures for "Melanoidins Suppress Cucumber and Tomato Diseases Incited by Pathogens that are Oomycetes, Bacteria and a Virus"

**Supplements**

Damping off (%)

| Time (Days) | 0 | 0.03 | 0.1 |
| --- | --- | --- | --- |
| 7 | 0 | 0 | 0 |
| 10 | 24.8 | 8.16667 | 8.2 |
| 12 | 24.8 | 8.33333 | 16.6 |
| 14 | 41.2 | 16.5 | 16.6 |
| 17 | 41.2 | 16.5 | 24.8 |

**Fig. S1** Effect of M on Oomycetes incited soilborne disease in cucumber – Pythium damping off. M spray was applied at concentrations of 0.03 and 0.1% on the day of seedlings transplanting in the *Pythium aphanidermatum* amended growth mixture and 7 days afterwards. Damping off was evaluated every 2 to 3 days and is presented as percent of all plants in each pot. Percent data at each evaluation date followed by a different letter are significantly different according to one-way ANOVA with Tukey’s HSD test. Default significance levels were set at α = 0.05. Bars = SEs.

Severity (%)

|  | 9d |
| --- | --- |
| Control | 6.49 |
| M 0.1% spray | 1.925 |
| M 0.1% drench | 1.745 |

**Fig. S2** Effect of M on an Oomycetes incited foliar disease in cucumber – Control of *Pseudoperonospora cubensis* cucumber downy mildew (PcCDM) by spraying and drenching 0.1% M. M was applied at days 0 and 3 and the severity of disease was evaluated on day 9. Severity of PcCDM was evaluated on a 0 to 100% scale, in which 0 = healthy leaves and 100% = leaves completely covered by the damage symptoms. Values followed by a different letter are significantly different according to one-way ANOVA with Tukey’s HSD test. Default significance levels were set at α = 0.05. Bars = SEs.

Disease severity (%)

| Time since 1st application (days) | M 0.1% | Control |
| --- | --- | --- |
| 1 | 0 | 0 |
| 5 | 2.8 | 6 |
| 11 | 3.6 | 7.4 |
| 12 | 5.6 | 12.4 |
| 13 | 11 | 19.4 |

**Fig. S3** Effect of M on late blight caused by *Phytophthora infestans* in tomato plants. Plants were treated by 0.1% M spray at days at days 0 and 5 and disease severity was evaluated until day 13 after the initiation of M applications. Severity of late blight was evaluated on a 0 to 100% scale, in which 0 = healthy leaves and 100% = leaves and stems completely covered by the damage symptoms. Values in each date followed by a different letter are significantly different according to one-way ANOVA with Tukey’s HSD test. Default significance levels were set at α = 0.05. Bars = SEs.

Disease severity (%)

| Time after planting (days) | Control | M 0.03% spray | M 0.03% drench |
| --- | --- | --- | --- |
| 1 | 0 | 0 | 0 |
| 3 | 4.3 | 4.6 | 5.4 |
| 6 | 14 | 4.8 | 6.2 |
| 10 | 17 | 5.3 | 7 |
| 13 | 52.5 | 19 | 24.7 |

**Fig. S4** Effect of M on the severity of *Pseudomonas syringae* pv*. tomato* bacterial speck (PstBS) naturally developed on leaves of tomato plants. 0.03% M was sprayed and drenched at 0 and 4 days after experiment initiation and evaluated through 13 days. Severity of PstBS was evaluated on a 0 to 100% scale, in which 0 = healthy leaves and 100% = leaves completely covered by the damage symptoms. Values in each date followed by a different letter are significantly different according to one-way ANOVA with Tukey’s HSD test. Default significance levels were set at α = 0.05. Bars = SEs.
